# A silk-based platform to stabilize phenylalanine ammonia-lyase (PAL) for orally administered enzyme replacement therapy

**DOI:** 10.1101/2022.04.14.488340

**Authors:** Luciana d’Amone, Vikas D. Trivedi, Nikhil U. Nair, Fiorenzo G. Omenetto

## Abstract

Phenylalanine ammonia-lyase (PAL) has gained attention in recent years for the treatment of phenylketonuria (PKU), a genetic disorder that affects ∼1 in 15,000 individuals globally. However, the enzyme is easily degraded by proteases, unstable at room temperature, and is currently administered in PKU patients as daily subcutaneous injections. We report here the stabilization of the PAL from *Anabaena variabilis*, which is currently used to formulate pegvaliase, through incorporation in a silk fibroin matrix. The combination with silk stabilizes PAL at 37 °C. In addition, in vitro studies showed that inclusion in a silk matrix preserves the biological activity of the enzyme in simulated intestinal fluid, enabling the oral administration of pegvaliase for the treatment of PKU.

## INTRODUCTION

Phenylketonuria (PKU) is an autosomal recessive disorder of amino acid metabolism that affects 0.45 million individuals globally.^1^ The indication presents as partial or complete loss in phenylalanine hydrolase (PAH) activity, causing elevated blood concentrations of phenylalanine and its aberrant metabolic phenylketone byproducts. If untreated, the disorder results in developmental delay or severe irreversible intellectual disability, motor deficit, and seizure. ^2^

Phenylalanine ammonia lyase (PAL) is the active ingredient in the only FDA-approved enzyme for enzyme replacement therapy to treat classical PKU – pegvaliase/Palynziq – as well as in several investigational therapeutics, including an engineered probiotic.^1^ PAL degrades phenylalanine to ammonium and *trans*-cinnamic acid, which is then excreted in the urine as hippuric acid, thus reducing the systemic phenylalanine concentration.^3^ PAL is currently available in pegylated form, to minimize immunogenicity and improve pharmacokinetics, and is self-administered daily as up to three subcutaneous injections.^4^ However, patient compliance is a major issue due to requirements of daily injections and potential for adverse events, such as injection site reactions or anaphylactic shock.^3^

The use of orally administered PAL ^1^ would decrease the risk of immune reactions and improve patient compliance, quality of life, and outcomes. Yet, the development of such PAL enzyme replacement therapies faces obstacles because of the PAL enzyme instability at room temperature and its rapid inactivation through denaturation or proteolytic degradation in the gut. ^5-8^

We present here a silk fibroin-based drug delivery system to address the stability limitations of PAL. Regenerated silk fibroin, a natural protein derived from the cocoons of *Bombyx mori* silkworms is biocompatible, biodegradable, and can be used to stabilize biomolecules, which makes it a good carrier for intrinsically labile molecules such as PAL. ^9-12^ In this study, we show that silk fibroin can preserve intact bioactivity of PAL at room temperature when it is embedded in solid film formats. These films allowed for the stabilization of the enzyme for over a month without the need for refrigeration. Furthermore, *in vitro* studies revealed that the silk film enables controlled release of PAL in simulated intestinal fluid over 20 h, resulting in a promising oral delivery option for the treatment of PKU (**Figure 1**).

**Figure 1.**
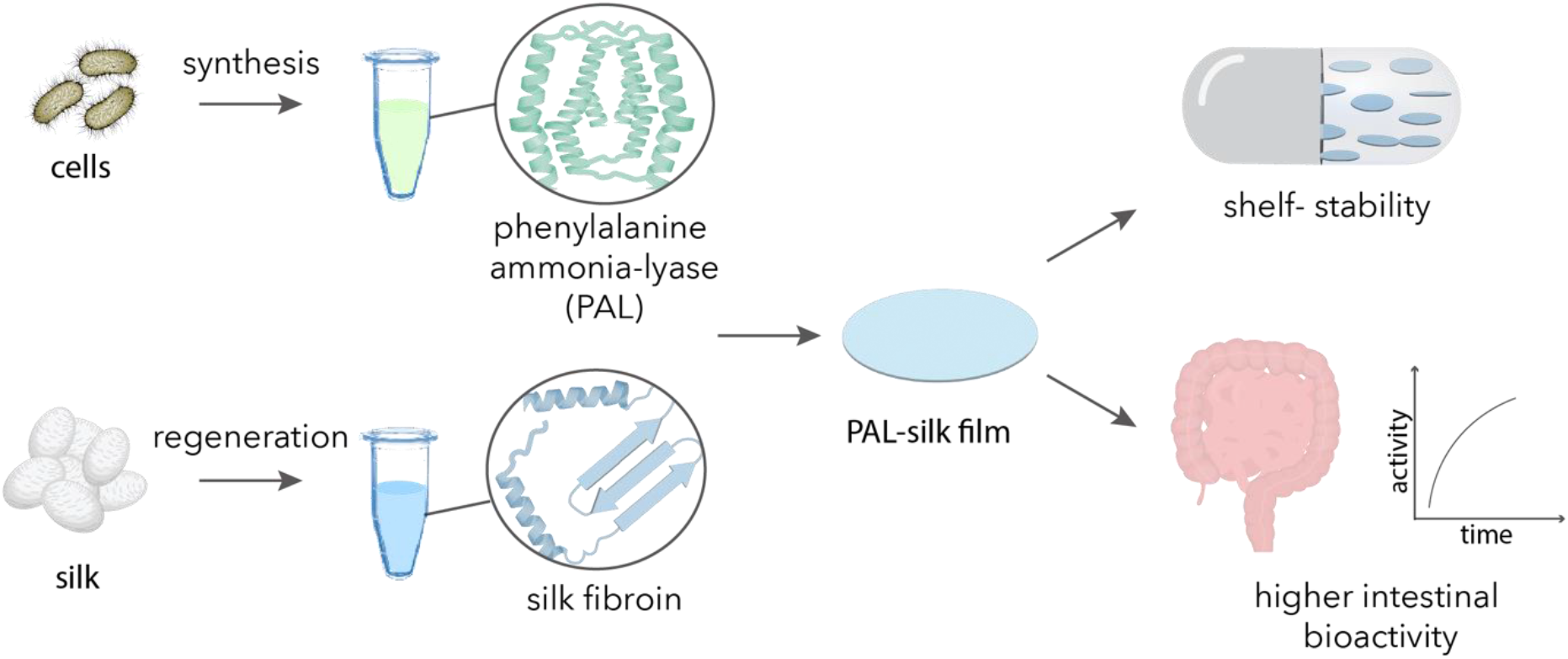
Conceptual overview of this work. Phenylalanine ammonia-lyase (PAL) expressed in recombinant *E. coli* cells and silk fibroin extracted from *B. mori* cocoons are combined to prepare shelf-stable drug delivery systems that preserve the biological activity of the therapeutic enzyme in intestinal fluids.

## MATERIAL AND METHODS

### Reagents

Sodium carbonate, lithium bromide, and phosphate buffered saline (PBS) were purchased from Sigma-Aldrich (St. Louis, MO). Silk cocoons from *Bombyx mori* silkworm were purchased from Tajima Shoji Co. (Yokohama, Japan). Deionized (DI) water with a resistivity of 18.2 MΩ cm was obtained with a Milli-Q reagent-grade water system and used to prepare aqueous solutions. Phenylalanine powder was purchased from Tokyo Chemical Industry (Portland, OR). Artificial gastric fluid (pH 1.5) and simulated salivary fluid (pH 6.8) were purchased from Biochemazone (Alberta, Canada), and simulated intestinal fluid (pH 7.4–7.6) was purchased from Ricca Chemical Company (Arlington, TX). Pancreatin was purchased from Thermo Fisher Scientific (Waltham, MA).

### Heterologous expression and purification of PAL

The previously engineered high activity quadruple mutant T102E-M222L-C503S-C565S *Anabaena variabilis* PAL ^1^ was expressed using pBAV1K (Cm^R^) under constitutive T5 promoter in *E. coli rph*^+^. The strain was cultured in lysogeny broth (LB) (VWR International, Randor, PA) at 37 ºC with rotary shaking at 250 rpm. For purification 50 mL of cells were cultured, pelleted and processed for purification as described earlier.^13^ Briefly, the cell pellet was weighed, resuspended (1:5 w/v) in PBS and lysed on ice via sonication (10 cycles of 30 s ON and 1.5 min OFF, output 40 %). The cell homogenate was centrifuged at 20,000 × g for 10 min and the clear supernatant was loaded on to 2 mL Ni-NTA matrix. The protein-bound matrix was washed with 5 column volumes of PBS containing 20 mM imidazole. The enzyme was eluted using 500 mM imidazole in PBS. The eluted enzyme was dialyzed twice (1:100 v/v) against PBS and stored at 4 °C until further processing and analysis. The purified enzyme was analyzed using SDS-PAGE and assayed as described previously. ^14^

### Silk fibroin extraction

Regenerated silk fibroin solution was extracted from *Bombyx mori* cocoons as previously reported^15^. Briefly, silk cocoons were shredded, separated from impurities, and boiled in a 20 mM sodium carbonate solution to remove sericin. The fibers were rinsed with DI water three times for 20 minutes and air-dried overnight. The dried silk mats were subsequently dissolved in a 9.3 M lithium bromide at 60 °C for 4 h. The chaotropic salt was then removed through dialysis (dialysis membranes MWCO 35 kDa) against DI water for 48 h, changing the water 6 times at regular intervals, which yielded a 7-8 wt% silk solution. The final solution was centrifuged at ∼12,700 × g for 20 min, filtered to separate remaining impurities, and stored at 4 ºC.

### Preparation of silk films

7 wt% silk solution was combined with the dialyzed solution of PAL to prepare 2 wt% and 4 wt% silk films at 8 μg and 16 μg PAL/film. The mix was aliquoted and 80 μL were cast on Petri dishes and allowed to dry overnight at room temperature (∼20 ºC). The films were stored at room temperature before analysis or in an incubator (37 ºC) for stability studies.

### Characterization of silk films

Silk film secondary structural analysis was measured using Fourier Transform Infrared (FTIR) Spectroscopy (Bruker Invenio S diamond ATR). A measurement of 64 scans was collected for each sample at a resolution of 4 cm^-1^, which was acquired over a wavenumber range of 400–4000 cm^-1^. Spectral analysis was performed with OPUS (Bruker Optics, Inc.). Background absorption due to water was subtracted from the sample spectra to obtain a flat recording.

### PAL activity measurement

Enzyme activity was measured after film preparation to establish baseline activity (time, t = 0). 3 samples (N=3) were used for each point of analysis. Silk coated samples were dissolved in 80 μL of DI water and incubated for 2 min. Enzyme activity was then measured as described previously ^14^. Briefly, the 10 μL of dissolved films were mixed with 190 μL of pre-warmed phenylalanine (30 mM final concentration) in PBS in a 96-well F-bottom UVStar (Greiner Bio-One, Kremsmünster, Austria) microtiter plate. Absorbance at 290 nm was measured every 30 s at 37 ºC using a SpectraMax M3 (Molecular Devices) plate reader. The same protocol was used to analyze the enzyme activity in film stored at 37 ºC. Films stored at 37 ºC were dissolved and analyzed after 7, 14, and 42 days. The activity of PAL embedded in the silk film was compared with the activity of free PAL in solution stored at 4 ºC and 37 ºC.

### In vitro release in simulated fluids

To evaluate the performance of silk-coated PAL in simulated body fluids, artificial gastric fluid (Biochemazone, BZ175), salivary fluid (Biochemazone, BZ327), and intestinal fluid (Biochemazone, BZ176) containing pancreatin (Ricca R7109750-500A) were used as a proxy for in vivo environments. Silk-coated PAL or the equivalent amount of free enzyme were added to the fluids, incubated at 37 ºC, and the enzyme stability was assayed over the course of 24 h.

## RESULTS AND DISCUSSION

### Incorporation of PAL in silk films

Silk fibroin is biocompatible, biodegradable, and has excellent mechanical properties, does not cause an immune or a significant inflammatory reaction, which renders fibroin an excellent candidate in drug delivery systems.^16, 17^ To obtain water-soluble silk fibroin, silk cocoons were boiled through a degumming process used to remove the water-soluble protein glue, sericin, which bonds the fibroin together in the natural silk fiber. After degumming, silk fibers were dried in a cabinet overnight and dissolved to obtain a water-based fibroin solution. PAL was purified to apparent homogeneity using Ni-NTA affinity matrix (**Figure 2a**) and incorporated into silk solution on Petri dishes to generate the PAL-silk films. Since silk may exist in its amorphous water-soluble structure (silk I) or crystalline insoluble structure (silk II), FT-IR was used to analyze the structure of the dry films.^18^

**Figure 2.**
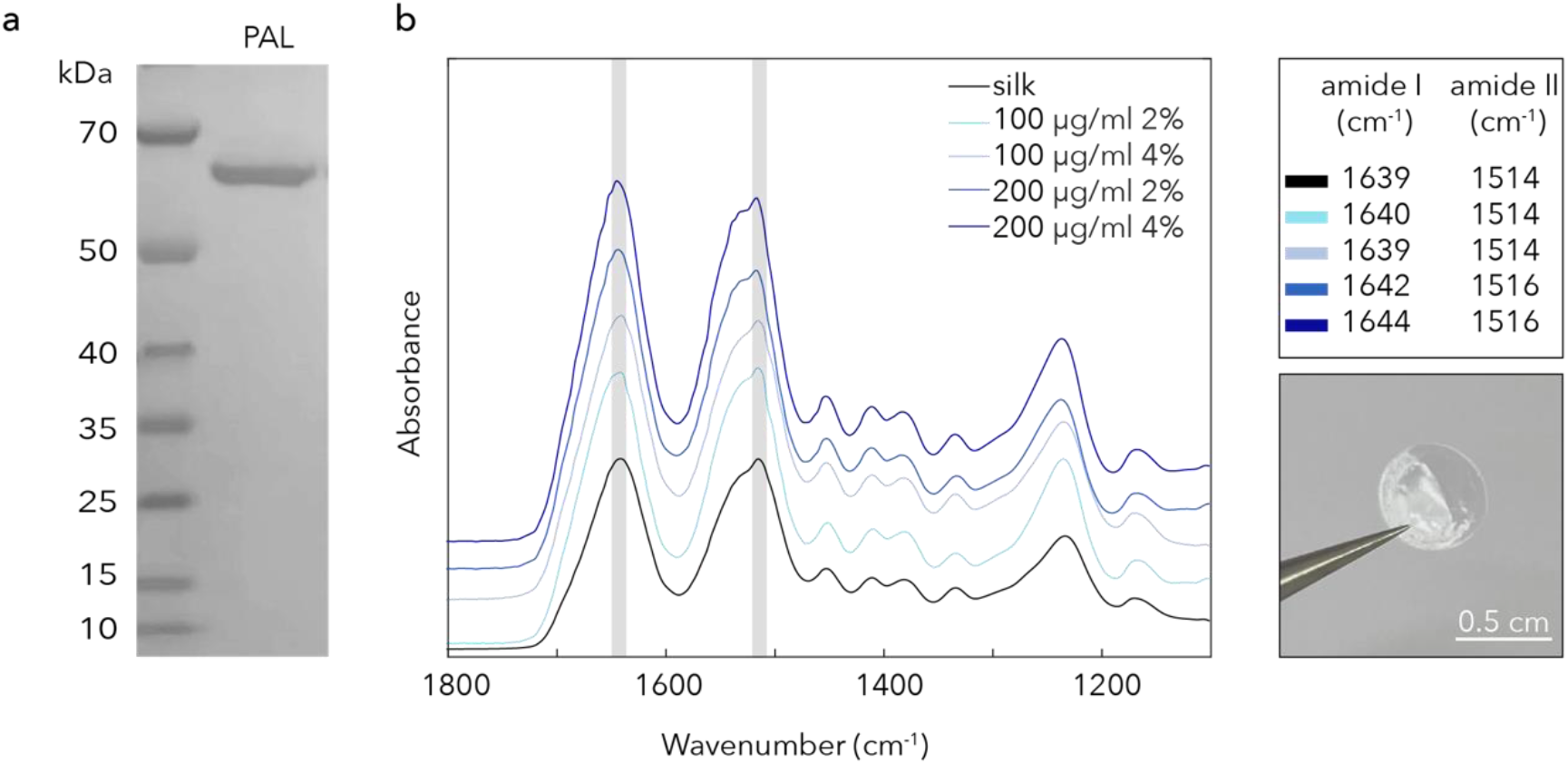
Structural characterization of the PAL-silk films. (a) SDS-PAGE confirms the molecular weight of PAL expressed from *E. coli* (b) FT-IR of silk films at different concentrations. Top-right insert reports the peak values of the amide I and II of silk fibroin in the different films, bottom shows a photograph of the PAL-silk films.

We tested four different combinations of silk fibroin and PAL, including two silk concentrations (2 and 4 wt%) and two of PAL (8 and 16 μg/film) to evaluate possible structural changes in the films. The 1700–1500 cm^-1^ infrared spectral region was used to evaluate the absorption of the peptide backbone of the silk amide I (1700–1600 cm^-1^) and amide II (1600–1500 cm^-1^), which are used for the analysis of the different secondary structure of silk fibroin (**Figure 2b**). ^18-20^ The assigned bands revealed the presence of the silk I structure, and no silk II or beta-sheet, indicating that the PAL-silk films were mostly in the amorphous structures of random coils and/or helix conformations.

### Shelf-stability of PAL in silk films

To evaluate possible changes in the bioactivity of PAL derived from inclusion in the silk matrix, the initial activity (i.e., t = 0) relative to equivalent amounts of free enzyme (**Figure 3a**) was evaluated. Both silk and PAL concentrations influenced the dried matrices bioactivity. Specifically, excess of either silk or PAL decreased overall bioactivity – demonstrated by low activity of both 100 μg/mL PAL 4 % silk and 200 μg/mL PAL in 2 % silk films. Conversely, when PAL loading was matched with that of silk (100 μg/mL PAL 2 % silk and 200 μg/mL PAL in 4 %), the system demonstrated activity equivalent to that of free enzyme suggesting that the 2 % silk films have insufficient solid-state silk matrix to stabilize high loading of PAL. Under these conditions, PAL may be aggregating, leading to a loss in bioactivity. Additional studies will be required to evaluate potential conformational changes of PAL in the silk matrix. Higher silk concentrations (4 %) may lead to stronger interactions between the proteins, decreasing the rate of PAL release, resulting in lower observed enzyme activity when loaded at 100 μg/mL. Increasing enzyme loading seems to overcome this issue, as shown by the higher enzymatic activity of the 200 μg/mL PAL with 4 % silk. Thus, with the right formulation, silk-embedded PAL retains the activity equivalent to that of the free enzyme.

**Figure 3.**
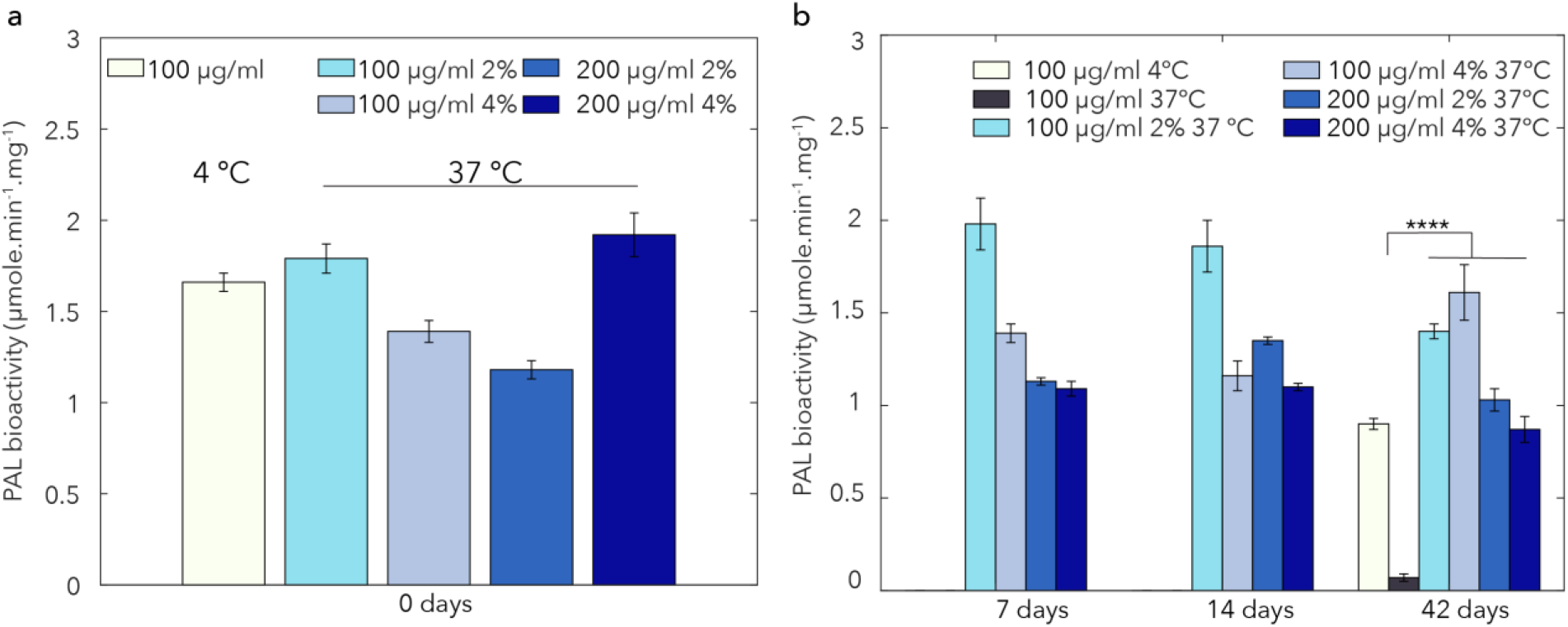
PAL stability in silk films. (a) Bioactivity of PAL in dried silk films at different concentrations of silk and PAL at time 0 (b) Bioactivity of the PAL-silk films after storage at 37 ºC for 7, 14, and 42 days. Values are average ± standard deviation of a minimum of N = 3 samples for each group. **** indicates significant difference between samples and controls (at 4 and 37ºC) (p< 0.0001)

Stabilization of PAL in silk for prolonged storage without refrigeration was also evaluated. Silk fibroin is inherently stable to moisture and changes in temperature thanks to its heterogeneous structure that generates protective microenvironments around labile molecules, including enzymes, drugs, and genomic content. ^9, 15, 21, 22^ PAL is currently on the market as a prefilled syringe that needs to be refrigerated at 2–8 ºC. To perform an accelerated aging test, silk-PAL films were stored at 37 ºC in an incubator and dissolved and analyzed after 7 days, 14 days, and 42 days. The bioactivity of PAL in the silk film at different concentrations was compared with the stability of the enzyme solution without the silk matrix stored at 4 ºC and 37 ºC. The stability studies showed that PAL-silk films at different concentrations preserved the bioactivity of the enzyme for over a month, with the 100 μg/mL 2 % films being the best performers (**Figure 3b**). After 42 days, the bioactivity of 100 μg/mL PAL in 2 % film was statistically higher than the activity of PAL in solution stored at 37 ºC, which had completely lost its activity (**Figure 3b**). Overall, 100 μg/mL PAL loaded in 2 % silk demonstrated both the most favorable initial reaction rate and long-term stability.

### Preservation of PAL by silk films in simulated digestive fluids

Changes in ionic strength and pH that proteins may experience in vivo after oral administration may destabilize them, resulting in a loss in activity. Furthermore, the presence of proteases in biological fluids can influence the degradation profile of proteins such as silk and PAL, affecting both the dissolution and release of the active molecule from the carrier and the bioactivity of the active ingredient. To evaluate the effectiveness of silk in preserving the activity of PAL after oral administration, the stability of the silk-based drug delivery system was evaluated in simulated biological fluids. Since the 100 μg/mL 2 % films were found to be the most promising during stability studies, these films were selected for further analysis and tested their stability in simulated salivary, gastric, and intestinal fluids and compared against the free enzyme. As an experimental control, we used PAL dissolved in PBS. Without silk, PAL lost almost all its activity after 2 h in simulated saliva (pH 6.8) and intestinal fluids (pH 7.5) and within minutes in simulated gastric fluid (pH 1.5) (**Figure 4a**). PAL encapsulated in the silk matrix was more stable after addition of simulated gastric fluid, but the activity of the enzyme decreased in the first 2 h. Visual inspection of the samples revealed that the PAL-silk conjugate precipitated after 4 h in salivary fluid, leading to loss in enzyme activity (**Figure 4b**). The 100 μg/mL 2 % films released PAL progressively in simulated intestinal fluid spiked with pancreatin, leading to a sustained release from the silk matrix over 20 h, as suggested by the progressively increasing activity over time. This makes silk films an excellent oral delivery optional for PAL with sustained activity by slow-release. Due to the instability in gastric fluids, PAL-silk films may need to be formulated in capsules that protect can protect the enzyme from gastric fluids (e.g., recombinant gelatin or cellulose acetate phthalate).

**Figure 4.**
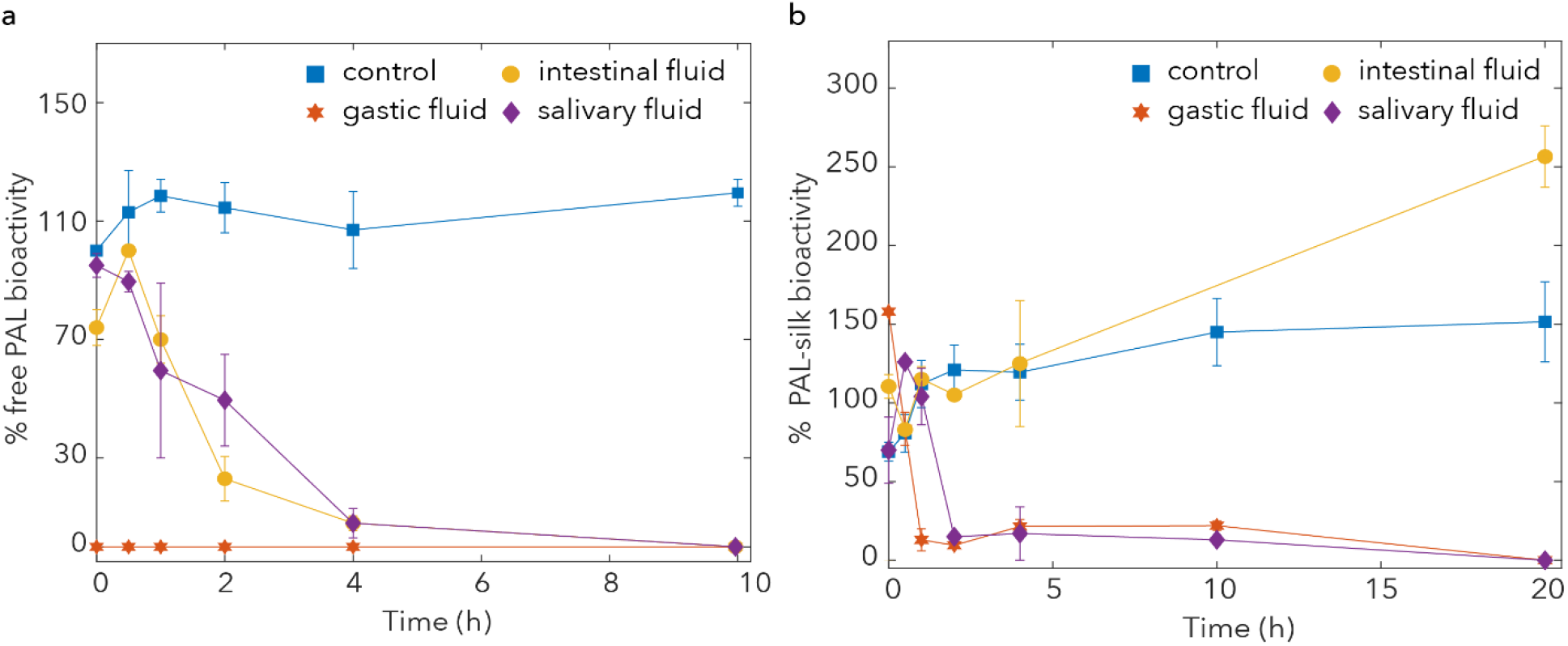
Stability of PAL-silk films in simulated digestive fluids. (a) Bioactivity of free PAL and (b) PAL in silk films after incubation in simulated salivary, gastric, and intestinal fluids. The control represents PAL dissolved in PBS only.

## CONCLUSIONS

The physical-chemical properties of silk fibroin underscore the utility of silk-based formats as promising drug delivery systems through a relatively simple and straightforward manufacturing process. Self-assembled silk fibroin processed into film formats resulted in a stable and highly effective delivery carrier for the enzyme PAL. These films preserved the stability of the enzyme at 37 ºC for over a month and enabled controlled release of the therapeutic enzyme in simulated intestinal fluid for 20 h. The ability of silk to stabilize labile molecules in simulated fluids presenting proteolytic enzymes supports new applications of silk fibroin for oral delivery of therapeutic proteins.

